# Activating the NFE2L1-ubiquitin-proteasome system by DDI2 protects from ferroptosis

**DOI:** 10.1101/2023.07.04.547652

**Authors:** Anahita Ofoghi, Stefan Kotschi, Imke L. Lemmer, Daniel T. Haas, Nienke Willemsen, Batoul Bayer, Sophie Möller, Stefanie Haberecht-Müller, Elke Krüger, Natalie Krahmer, Alexander Bartelt

**Affiliations:** Institute for Cardiovascular Prevention (IPEK), Faculty of Medicine, Ludwig-Maximilians-University, Munich, Germany; Institute for Diabetes and Cancer (IDC), Helmholtz Center Munich, German Research Center for Environmental Health, Neuherberg, Germany; Institute for Diabetes and Obesity (IDO), Helmholtz Center Munich, German Research Center for Environmental Health, Neuherberg, Germany; Institute of Medical Biochemistry and Molecular Biology, University Medicine Greifswald, Greifswald, Germany; German Center for Cardiovascular Research (DZHK), Partner Site Munich Heart Alliance, Munich, Germany; Department of Molecular Metabolism & Sabri Ülker Center for Metabolic Research, Harvard T.H. Chan School of Public Health, Boston, USA

**Keywords:** Ferroptosis, ubiquitin, proteasome, NFE2L1, DDI2

## Abstract

Ferroptosis is an iron-dependent, non-apoptotic form of cell death initiated by oxidative stress and lipid peroxidation. Recent evidence has linked ferroptosis to the action of the transcription factor Nuclear factor erythroid-2 derived,-like-1 (NFE2L1). NFE2L1 regulates proteasome abundance in an adaptive fashion, maintaining protein quality control to secure cellular homeostasis, but the regulation of NFE2L1 during ferroptosis and the role of the ubiquitin-proteasome system (UPS) herein are still unclear. In the present study, using an unbiased proteomic approach charting the specific ubiquitylation sites, we show that induction of ferroptosis leads to recalibration of the UPS. RSL3-induced ferroptosis inhibits proteasome activity and leads to global hyperubiquitylation, which is linked to NFE2L1 activation. As NFE2L1 resides in the endoplasmic reticulum tethered to the membrane, it undergoes complex posttranslational modification steps to become active and induce the expression of proteasome subunit genes. We show that proteolytic cleavage of NFE2L1 by the aspartyl protease DNA-damage inducible 1 homolog 2 (DDI2) is a critical step for the ferroptosis-induced feed-back loop of proteasome function. Cells lacking DDI2 cannot activate NFE2L1 in response to RSL3 and show global hyperubiquitylation. Genetic or chemical induction of ferroptosis in cells with a disrupted DDI2-NFE2L1 pathway diminishes proteasomal activity and promotes cell death. Also, treating cells with the clinical drug nelfinavir, which inhibits DDI2, sensitized cells to ferroptosis. In conclusion, our results provide new insight into the importance of the UPS in ferroptosis and highlight the role of the DDI2-NFE2L1 as a potential therapeutic target. Manipulating DDI2-NFE2L1 activity through chemical inhibition might help sensitizing cells to ferroptosis, thus enhancing existing cancer therapies.

**Graphical abstract:** 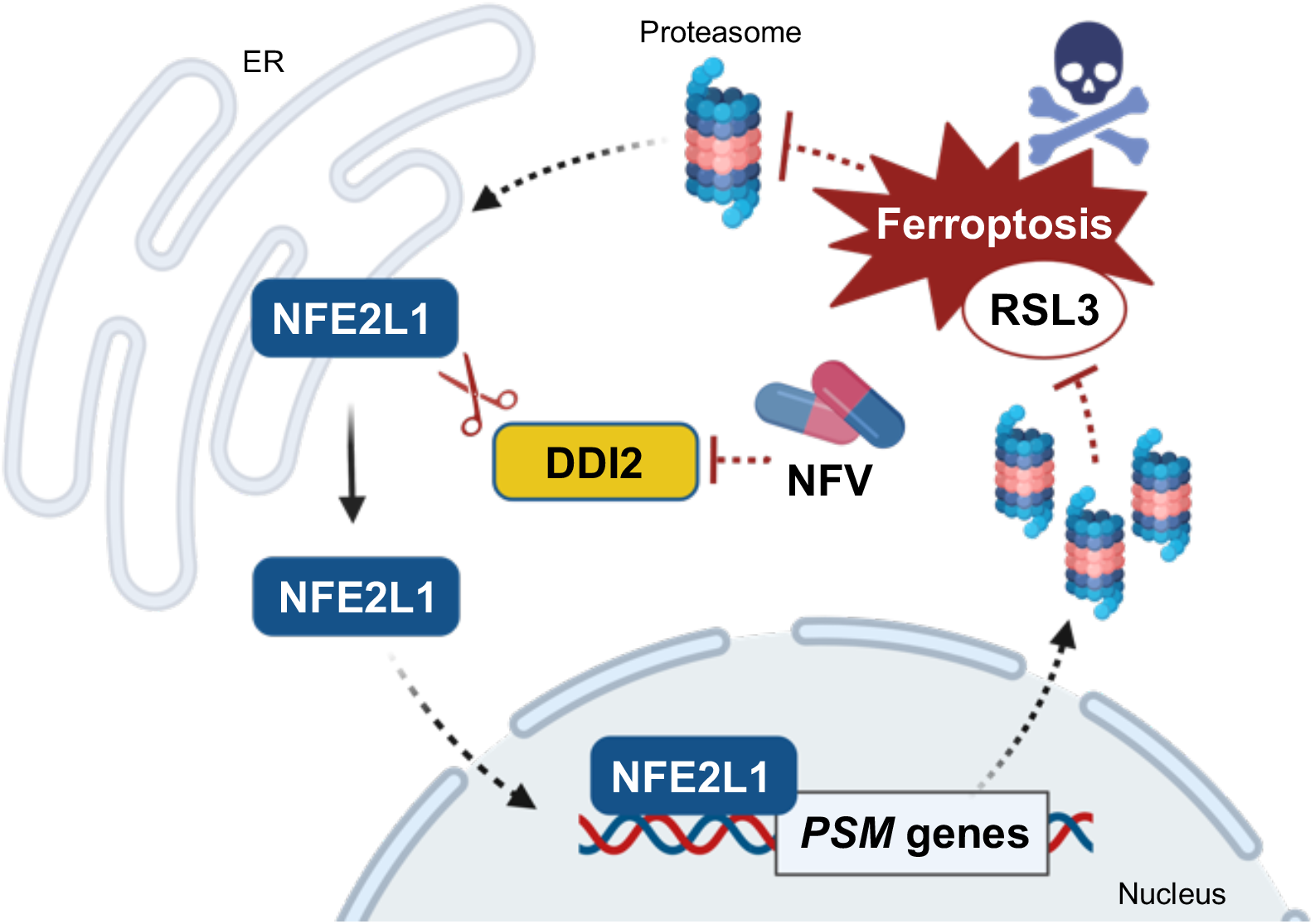

## Introduction

Regulated cell death is initiated by intracellular and extracellular perturbations that trigger tightly orchestrated molecular programs (1). Ferroptosis is a form of non-apoptotic cell death mediated by iron-dependent lipid peroxidation and loss of plasma membrane integrity (2, 3). Recent studies have implicated ferroptosis in several pathologies, such as neurodegeneration and cancer (4). Execution of ferroptosis is tightly linked to lipid and glutathione metabolism (5) and several chemical compounds have been shown to induce ferroptosis. Glutathione peroxidase 4 (GPX4) is a critical enzyme protecting from lipid reactive oxygen species (ROS) formation and, thus, from ferroptosis by most prominently uses glutathione for its antioxidative activity (6). While depletion of the reduced glutathione pool predisposes cells to ferroptosis, the compound RSL3 directly inhibits GPX4 (2, 5, 7). Thus, oxidative stress is sensed and mitigated by GPX4 and its inactivation leads to lipid peroxidation and cell death in cells and mouse models (8). Interestingly, ferroptosis is linked to adaptive changes in protein homeostasis, as ferroptosis initiation is associated with diminished proteasomal activity and restoration of proteasomal activity protects cells from ferroptotic cell death (9). The ubiquitin-proteasome system (UPS) manages the degradation of unwanted, obsolete, or damaged proteins (10). The UPS controls almost all cellular processes by precision proteolysis, enabled by specific ubiquitin ligases (11). Attachment of ubiquitin to these proteins is the prerequisite for the ATP-dependent degradation by the proteasome. The 26S proteasome is a multi-subunit protease complex, which consists of the barrel-shaped 20S proteolytic core particle capped with one or two 19S regulatory particles for binding to the polyubiquitylated proteins (12). While it is accepted that the UPS is a critical pillar of cellular health, the roles of UPS remodeling and the nature of these changes in ferroptosis remain unclear. We and others have shown that the transcription factor Nuclear factor erythroid-2, like-1 (NFE2L1, also known as TCF11 or NRF1) regulates ferroptosis (9, 13). NFE2L1 upregulates the expression of genes encoding proteasome subunit genes (14), thus restoring proteasomal activity and protecting from ferroptosis. NFE2L1 mediates the adaptive component of proteasomal activity and is subject to complex posttranslational regulation: NFE2L1 is a cap’n’collar transcription factor tethered in the endoplasmic reticulum membrane. In a simplified model, NFE2L1 undergoes deglycosylation by N-glycanase 1 (NGLY1, also known as PNGase) and proteolytic cleavage by the protease DNA damage-inducible 1 homolog 2 (DDI2) (15, 16). However, the resulting fragment containing the bZIP-DNA-binding domain-containing is virtually absent in most cells, as it is being continuously ubiquitylated and degraded by the proteasome (14, 17). Upon treatment with chemical proteasome inhibitors or if the levels of ubiquitylated proteins exceed the capacity of the available proteasomes, NFE2L1 escapes degradation and restores proteasomal activity(14). However, the impact of DDI2-mediated activation of NFE2L1 on UPS in ferroptosis remains unknown. Here, we determine the impact of ferroptosis on UPS and global ubiquitylation using an unbiased MS approach. Furthermore, we demonstrate a critical role of DDI2-mediated activation of the NFE2L1 pathway in calibrating the UPS for ferroptosis protection.

## Methods

### Cell culture, reverse transfection, and treatments

Human EA.hy926 cells were used for cellular experiments. DDI2 KO cells were created previously (18), and cultured in DMEM Glutamax (Thermo), supplemented with 10 % v/v fetal bovine serum (FBS, Sigma) and 1 % v/v Penicillin-Streptomycin (10.000 U/ml, Thermo). Cells were incubated at 37 °C, 5 % v/v CO_2_ and were passaged two to three times a week. For experiments 300,000, 100,000, and 25,000 cells were seeded at 6, 24, and 96-well cell culture plates, respectively. Primary GPX4 mutant fibroblasts (derived from a patient with a homozygous mutation c.647G>A in exon 6 of GPX4) and control cells were kindly provided by Sanath K. Ramesh (curegpx4.org). These cells were incubated in DMEM GlutaMax supplemented with 10 % v/v FBS and 1 % v/v PS, supplemented with 10 μM ferrostatin-1 (SelleckChem). Reverse transfection with SMARTpool siRNA (Dharmacon) was performed using 30 nM of siRNA in RNAiMAX (Thermo). Cells were incubated for 48 h after transfection, and then treatments were performed. For overexpression experiments, 250 ng of each plasmid construct (Empty vector, DDI2 WT, DDI D252, NFE2L1 WT, and NFE2L1-8ND) was added to TransIT-X2® Transfection Reagent (Mirus) and after 10 min incubation, cells were added to the medium. For treatments, different concentrations of RSL3 (SelleckChem), FIN56 (SelleckChem) and ferrostatin-1(SelleckChem) were used as indicated. Inhibition of proteasome and DDI2 was performed using bortezomib (SelleckChem) and nelfinavir mesylate (Sigma), respectively. Cell viability was assessed by AquaBluer (MultiTarget Pharmaceuticals), by changing the medium of cells to 1:100 of AquaBluer in phenol red-free DMEM GlutaMax, after 20 h of treatment. Cell plates were incubated for 4 h at 37 °C incubator and using a Spark Reader (Tecan) fluorescence was measured at emission of 590 nm with an excitation of 540 nm.

### Protein extraction and immunoblotting

Cells were lysed in RIPA buffer (150 mM NaCl (Merck), 5 mM EDTA (Merck), 50 mM Tris pH 8 (Merck), 1 % v/v IGEPAL CA-630 (Sigma), 0.5 % w/v sodium deoxycholate (Sigma Aldrich), 0.1% w/v SDS (Roth),1 mM protease inhibitors (Sigma)) for 3 min in TissueLyser II (30 Hz; Qiagen). Cell lysates were centrifuged for 15 min (4 °C, 21,000 *g*) and the concentration of proteins in supernatant was determined by Pierce BCA Protein Assay (Thermo) according to the manual. Proteins were denatured for 5 min at 95 °C in Bolt LDS Sample buffer with 5 % v/v 2-mercaptoethanol (Sigma). 10-30 μg of proteins were loaded in Bolt 4-12 % Bis-Tris gels (Thermo) followed by transferring onto a 0.2 mm PVDF membrane (Bio-Rad) at 25 V, 1.3 A for 7 min. Membranes were blocked in TBS-T (200 mM Tris (Merck), 1.36 mM NaCl (Merck), 0.1 % Tween-20 (Sigma)) containing 5 % w/v milk powder for 1 h at room temperature after staining in Ponceau-S (Sigma Aldrich). Incubation by primary antibodies (Supplementary Tab. 1) was performed overnight at 4 °C followed by washing membranes three times for 10 min with TBS-T and incubation with secondary antibodies for 1 h at room temperature. SuperSignal West Pico PLUS (Thermo) was used for developing blots in a ChemiDoc MP imager (Bio-Rad). All uncropped blots can be found in Supplementary figs. 1-3.

### Proteasome activity

Cells lysis was performed in lysis buffer (40 mM Tris pH 7.2 (Merck), 50 nM NaCl (Merck), 5 mM MgCl2(6H2O) (Merck), 10 % v/v glycerol (Sigma), 2 mM ATP (Sigma), 2 mM 2-mercaptoethanol (Sigma). Proteasome Activity Fluorometric Assay II kit (UBPBio, J41110) was used to measure proteasome activity. BCA Protein Assay (Bio-Rad) was used to normalize the results to protein levels.

### Native PAGE

Cells were lysed in lysis buffer (50 mM Tris/HCl pH 7.5, 2 mM DTT, 10 % v/v glycerol, 5 mM MgCl2, 0.05 % v/v Digitonin, 2 mM ATP) containing phosphatase inhibitor (PhosphoStop, Roche Diagnostics) as described previously (19). Samples were incubated on ice for 20 min and centrifuged twice. Concentration of proteins was determined with Bio-Rad Protein Assay Kit II. 15 μg of protein were loaded in NuPAGE 3-8 % Tris-Acetate gels (Thermo) and run at constant voltage of 150 V for 4 h. Gels were kept for 30 min at 37 °C in activity buffer (1 mM MgCl2, 50 mM Tris, 1 mM DTT) with 0.05 mM substrate Suc-Leu-Leu-Val-Tyr-AMC (Bachem). ChemiDoc MP (Bio-Rad) was used to measure the fluorescent signal. Afterwards, to prepare samples for blotting gel was incubated in a solubilization buffer (2 % w/v SDS, 1.5 % v/v 2-Mercaptoethanol, 66 mM Na_2_CO_3_) for 15 min. Samples were transferred to a PVDF membrane at 40 mA through tank transfer. The membrane was kept for 1 h in the ROTI-block (Roth) and overnight in primary antibody (1:1000). The day after, the membrane was incubated for 3 h in the secondary antibody (1:10,000) at room temperature and developed as described above.

### NFE2L1 reporter assay

HEK293a cells stably expressing short half-life firefly luciferase driven by upstream activator sequence (UAS) promoter and a chimeric NFE2L1 in which the DNA-binding domain was replaced by the UAS-targeting Gal4 DNA-binding domain (20). The assay measures nuclear translocation and its transactivation by binding of NFE2L1-UAS to a luciferase promoter (20). 30,000 cells were seeded in 96-well plates and after treatment with RSL3, cells were lysed and luciferase emission was measured using Dual-Glo Luciferase Assay System (Promega) according to the manufacturer’s instructions.

### Protein digestion and purification for MS

Protein digestion and DiGly peptide enrichment was performed as described previously (21, 22). Cells were lysed in SDC buffer (1 % v/v SDC in 100 mM Tris-HCl, pH 8.5) followed by boiling for 5 min at 95 °C while shaking at 1000 rpm. Protein concentrations of lysates were determined after 15 min of sonication (Bioruptor, Diagenode, cycles of 30 s) using the Pierce BCA Protein Assay (Thermo). CAA and TCEP (final concentrations: 40 mM and 10 mM respectively) were added to 5 mg protein. After 10 min incubation of samples at 45 °C in the dark shaking at 1000 rpm, Trypsin (1:50 w/w) and LysC (1:50 w/w) were added. Samples then were kept overnight at 37 °C while shaking at 1000 rpm for digestion. For proteome analysis, sample aliquots (∼15 μg) were desalted in SDB-RPS StageTips (Empore). Briefly, samples were diluted with 1 % TFA in isopropanol to a final volume of 200 μl, loaded onto StageTips, and sequentially washed with 200 μl of 1 % v/v TFA in isopropanol and 200 μl 0.2 % v/v TFA/2 % v/v ACN. Peptides were eluted with freshly prepared 60 μl of 1.25 % v/v ammonium hydroxide (NH_4_OH)/80 % v/v ACN and dried using a SpeedVac centrifuge (Eppendorf). Dried peptides were resuspended in 6 μL buffer A (2 % v/v ACN/0.1 % v/v TFA). Di-Gly enrichment samples were diluted with 1 % v/v TFA in isopropanol (1:5). For peptide cleanup, SDB-RPS cartridges (Strata™-X-C, 200 mg/6 ml, Phenomenex Inc.) were equilibrated with 8 bed volumes (BV) of 30 % v/v MeOH/1 % v/v TFA and washed with 8 BV of 0.2 % v/v TFA. Samples were loaded by gravity flow and sequentially washed twice with 8 BV 1 % TFA in isopropanol and once with 8 BV 0.2 % v/v TFA/2 % v/v ACN. Peptides were eluted with 2 × 4 BV 1.25 % v/v NH4OH/80 % v/v ACN and diluted with ddH2O to a final of 35 % v/v ACN. Samples were dried via a SpeedVac Centrifuge overnight.

### DiGly peptide enrichment

We used the PTMScan® Ubiquitin Remnant Motif (Cell Signaling). The peptides were resuspended in 500 μL immunoaffinity purification (IAP) buffer and sonicated (Bioruptor) for 15 min. BCA Protein Assay (Thermo) was used to determine protein concentration. The antibody-coupled beads were cross-linked as previously described (22). One vial of antibody coupled beads was washed with cold cross-linking buffer (2000 *g*, 1 min). Subsequently the beads were incubated in 1 mL cross-linking buffer at room temperature for 30 min under gentle agitation. The reaction was stopped by washing twice with 1 mL cold quenching buffer (200 mM ethanolamine, pH 8.0) and incubating for 2 h in quenching buffer at room temperature under gentle agitation. Cross-linked beads were washed three times with 1 mL cold IAP buffer and directly used for peptide enrichment. For DiGly enrichment, 3 mg of peptide is used with 1/8 of a vial of cross-linked antibody beads. Peptides are added to the cross-linked beads and the volume is adjusted to 1 mL with IAP buffer and incubated for 2 h at 4 °C under gentle agitation. The beads are washed twice with cold IAP buffer and twice ddH_2_O via centrifugation. The enriched peptides were eluted by adding 200 μL 0.2 % v/v TFA onto the beads, incubating 5 min at 1400 rpm and centrifuging for 1 min at 100 *g*. The supernatant was then transferred to SDB-RPS StageTips and the peptides washed, eluted and dried as previously described for total proteome samples.

### LC-MS/MS proteome and ubiquitome measurements

Liquid chromatography of total proteome and ubiquitome samples was performed on an EASYnLCTM 1200 (Thermo) with a constant flow rate of 10 μL/min and a binary buffer system consisting of buffer A (0.1 % v/v formic acid) and buffer B (80 % v/v acetonitrile, 0.1 % v/v formic acid) at 60 °C. The column was in-house made, 50 cm long with 75 μm inner diameter and packed with C18 ReproSil (Dr. Maisch GmbH, 1.9 μm). The elution gradient for the ubiquitome started at 5 % buffer B, increasing to 25 % after 73 min, 50 % after 105 min and 95 % after 110 min. The gradient for the proteome started at 5 % buffer B and increased to 20 % after 30 min, further increased at a rate of 1 % per minute to 29 %, following up to 45 % after 45 min and to 95 % after 50 min. The MS was performed on a Exploris480 with injection of 500 ng peptide (Thermo). Fragmented ions were analyzed in Data Independent Acquisition (DIA) mode with 66 isolation windows of variable sizes. The scan range was 300 – 1650 m/z with a scan time of 120 min, an Orbitrap resolution of 120,000 and a maximum injection time of 54 ms. MS2 scans were performed with a higher-energy collisional dissociation (HCD) of 30 % at a resolution of 15,000 and a maximum injection time of 22 ms. The MS measurement of the proteome was performed equivalently, yet including HighField Asymmetric Waveform Ion Mobility (FAIMS) with a correction voltage of −50 V and a scan time of 60 min. The injection time for the full scan was 45 s and the MS2 injection time was set to 22 s.

### Proteome and Ubiquitome data analysis

DIA raw files were processed using Spectronaut (13.12.200217.43655) in directDIA mode. The FASTA files used for the search were: Uniprot Homo sapiens (29.03.2022) with 20609 entries, Uniprot Homo sapiens isoforms (29.03.2022) with 77157 entries and MaxQuant Contaminants for filtering: 245 entries. Analysis was performed via Perseus (version 1.6.2.3). For the ubiquitome samples the output was converted with the plugin “Peptide Collapse” (version 1.4.2). Values were log2-transformed and missing values were replaced by imputation from normal distribution with a width of 0.3 and downshift 1.8 separately for each sample. Ubiquitome was normalized to total proteome. Comparison between conditions of proteome and ubiquitome was performed via Student’s T-test in R 4.2.2 (*P* value cutoff 0.5).

### Gene expression analysis

RNA extraction was performed using Nucleospin RNA kit (Macherey Nagel), based on manufacturer’s instruction, and the concentration of RNAs were measured with a NanoDrop spectrophotometer (Implen). To prepare complementary DNA (cDNA), 500 ng RNA were added to 2 μL of Maxima™ H Master Mix 5x (Thermo Fischer) and the total volume was adjusted to 10 μL with H_2_O. The cDNA was diluted 1:40 in H_2_O, and 4 μL of cDNA, 5 μL of PowerUp™ SYBR Green Master Mix (Thermo), and 1 μL OF 5 μM primer stock (Supplementary Tab. 2) were used to measure Relative gene expression. Cycle thresholds (Ct) of gene of interest were measured using a Quant-Studio 5 RealTime PCR system (Thermo). Relative gene expression was normalized to TATA-box binding protein (*TBP*) levels by the ddCt-method.

### Statistics

All data were analyzed with Microsoft Excel, GraphPad Prism, and R (4.2.2). Data were visualized in GraphPad Prism and shown as mean ± standard deviation (SD). 1-way ANOVA with Dunnett’s Post-hoc Test was used when comparing three or more groups and one variable, and 2-way ANOVA followed by Tukey’s Test was used for comparing two groups with two variables. *P*-values lower than 0.05 were considered significant and are as such indicated in the graphs with an asterisk between groups.

## Results

### RSL3-induced ferroptosis diminishes proteasomal activity and activates NFE2L1

It remains largely unclear how the UPS is involved in the execution of ferroptosis. For two specific reasons this is important to understand: First, it is unclear if the regulation of UPS is a ferroptosis-specific event or simply a consequence of proteasome dysfunction. Second, as critical regulators of ferroptosis such as GPX4 have been shown to be regulated by UPS (9), understanding global and site-specific ubiquitylation will potentially aid the discovery of novel key players in ferroptosis. Treatment of EA.hy926 cells with the ferroptosis inducer RSL3 led to higher levels of ubiquitin (Fig. 1a and Supplementary fig. 4). Interestingly, the effect of RSL3 on ubiquitin levels was somewhat comparable if not higher to treatment of cells with the established chemical proteasome inhibitor bortezomib (BTZ) (Fig. 1a). We next asked if this effect also triggered the activation of NFE2L1, which an important part of UPS defense against inhibition of the proteasome. Indeed, treatment with RSL3 led to higher protein levels of the cleaved fragment of NFE2L1 (ca. 95 kDa) and lower levels of the full-length protein in a time- and dose-dependent fashion (Fig. 1b). This increase in cleaved fragment was associated with higher levels of NFE2L1 in the nucleus (Fig. 1c). Based on native PAGE analysis and measuring degradation of fluorogenic peptides, RSL3 diminished proteasomal activity (Fig. 1d,e). Hence, we tested the possibility that RSL3 may have off-target effects directly on proteasomal activity. At concentrations well-above what is used in our cell assays there was no direct inhibition of proteasomal activity in cell lysates, unlike what is seen with regular proteasome inhibitors. (Fig. 1f). This diminishes the possibility that the RSL3-induced decrease in proteasome activity and increase in ubiquitin levels are consequences of off-target pharmacological inhibition of the proteasome by direct binding of RSL3.

**Fig. 1:**
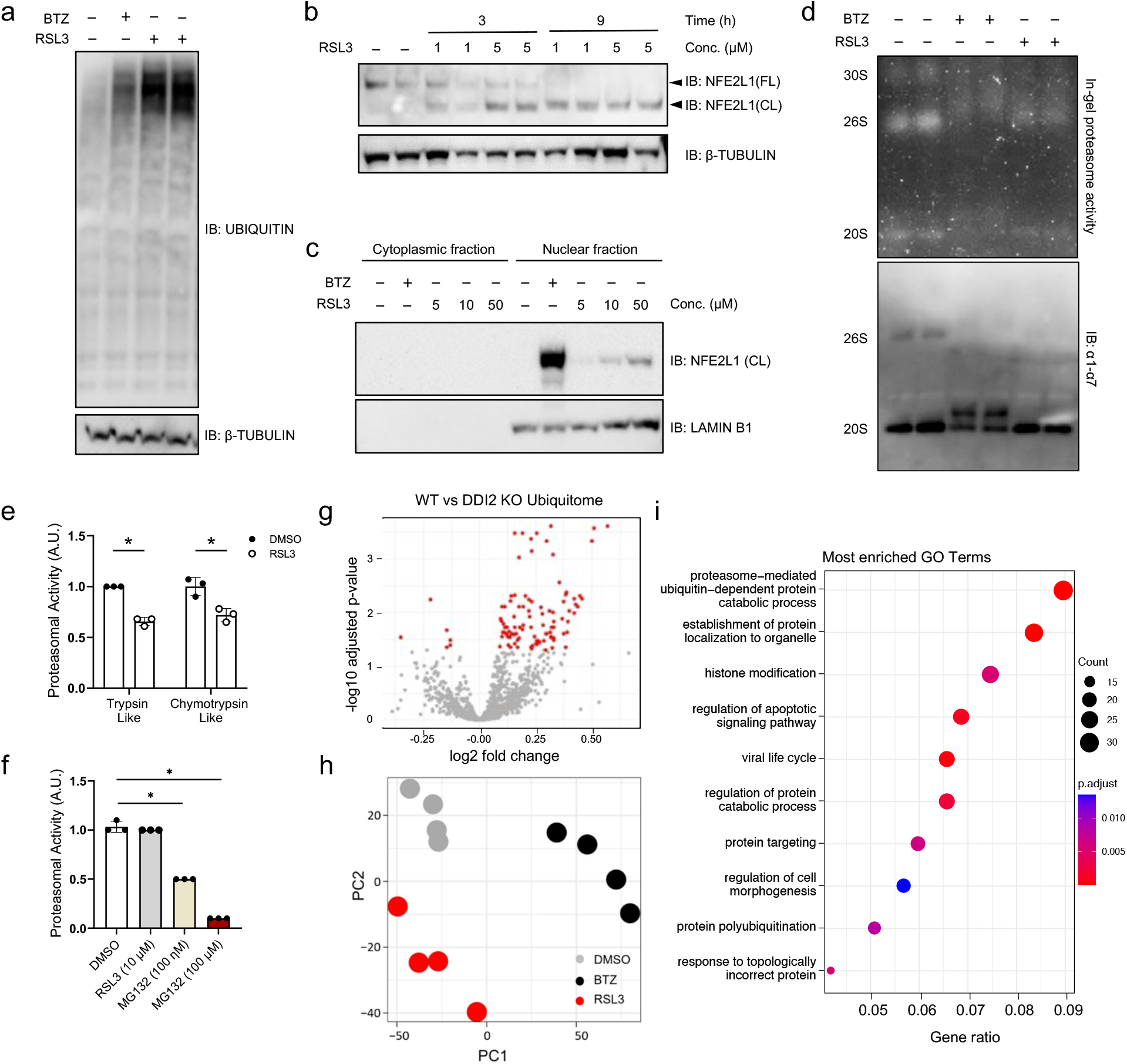
RSL3-induced calibration the UPS is distinct from proteasome inhibition. (a) Immunoblot of ubiquitin from EA.hy926 cells treated with 5 μM RSL3 and 100 nM bortezomib (BTZ) for 9 h and 3 h, respectively. (b) Immunoblot of NFE2L1 in EA.hy926 cells treated with indicated time points and concentrations of RSL3 (FL: full-length form ca. 120 kD; CL: cleaved ca. 95 kD). (c) Immunoblot of NFE2L1 in nuclear and cytoplasmic fractions isolated from EA.hy926 cells treated as indicated for 3 h (d) Native page of EA.hy926 cells treated with 5 μM RSL3 and 100 nM BTZ for 6 h with in-gel activity and immunoblot of the α1-7 (20S) subunits. (e) Proteasomal activity in EA.hy926 cells treated with 5 μM RSL3 for 3 h. (f) Proteasomal activity assay in EA.hy926 cells with the extracts being incubated with indicated concentrations and time points of RSL3 and proteasome inhibitor MG132. (g) Volcano Plot of the ubiquitome of EA.hy926 cells treated for 9 h with 5 μM RSL3 *P*_adj_ < 0.05 indicated in red. (n = 4 replicates) (h) Principal component analysis (PCA) of the ubiquitome of EA.hy926 cells treated with 5 μM RSL3 and 100 nM BTZ (n = 4 replicates). (i) Top 10 enriched gene ontology (GO) terms in the ubiquitome of EA.hy926 cells treated for 9 h with 5 μM RSL3.

### Remodeling of the ubiquitome in ferroptosis is distinct from proteasome inhibition

To better understand the calibration of UPS during ferroptosis, unequivocally determine the levels of ubiquitin, and to chart the specific lysine residues modified with ubiquitin, we performed a mass spectrometry-based analysis of the proteome and ubiquitylated proteins using the so-called Di-Gly method of immunoprecipitating ubiquitin remnant motifs (23). We identified 7063 peptides in the proteome and 3974 peptides in the ubiquitome. These analyses confirmed the increase in ubiquitin levels upon RSL3 treatment (Fig. 1g). To delineate and compare how RSL3 and BTZ treatments impact the ubiquitome, we performed principal component analysis and found that the three groups cluster significantly apart from each other (Fig. 1h). This indicates that the nature of ubiquitome remodeling induced by RSL3 is distinct from common proteasome inhibition, e.g. by BTZ. Next, to better understand the mechanisms involved in the execution of ferroptosis, we analyzed the cellular pathways implicated in the remodeling of the ubiquitome. Here, we found that gene ontology (GO) terms linked to UPS, cell cycle, cell death were among the differentially regulated pathways (Fig. 1i). In summary, this set of data highlights the remodeling of the ubiquitome, implicates the regulation of cell death signaling pathways by the UPS, and shows the activation of the proteostatic bounce-back response of NFE2L1 upon impairment of proteasomal activity upon RSL3-induced ferroptosis.

### DDI2 facilitates the RSL3-induced proteolytic activation of NFE2L1

As we observed an RSL3-induced increase in ubiquitin levels and NFE2L1 activation, we addressed the importance of the protease DDI2 for NFE2L1 cleavage under these conditions. We performed the next experiments in a cell line, in which DDI2 was deleted by CRISPR/Cas9 technology. Similar to BTZ treatment, RSL3 led to NFE2L1 activation in the parental cell line. However, in the DDI2 KO cells, neither BTZ nor RSL3 treatment led to changes in NFE2L1 protein levels (Fig. 2a), indicating that the activation of NFE2L1 during ferroptosis is critically dependent on DDI2. This effect on Nfe2l1 was also reproduced using FIN56 for ferroptosis induction (Supplementary fig. 4). To rule out that this is a cell line-dependent false negative effect, we re-expressed DDI2 in KO cells, and this restored BTZ-induced activation of NFE2L1 (Fig. 2b). Furthermore, reconstitution of DDI2 KO cells with a construct carrying proteolytically inactive DDI2 (DDI2-D252) did not restore the activation of NFE2L1 compared to re-expression of the WT protein (Fig. 2b).

**Fig. 2:**
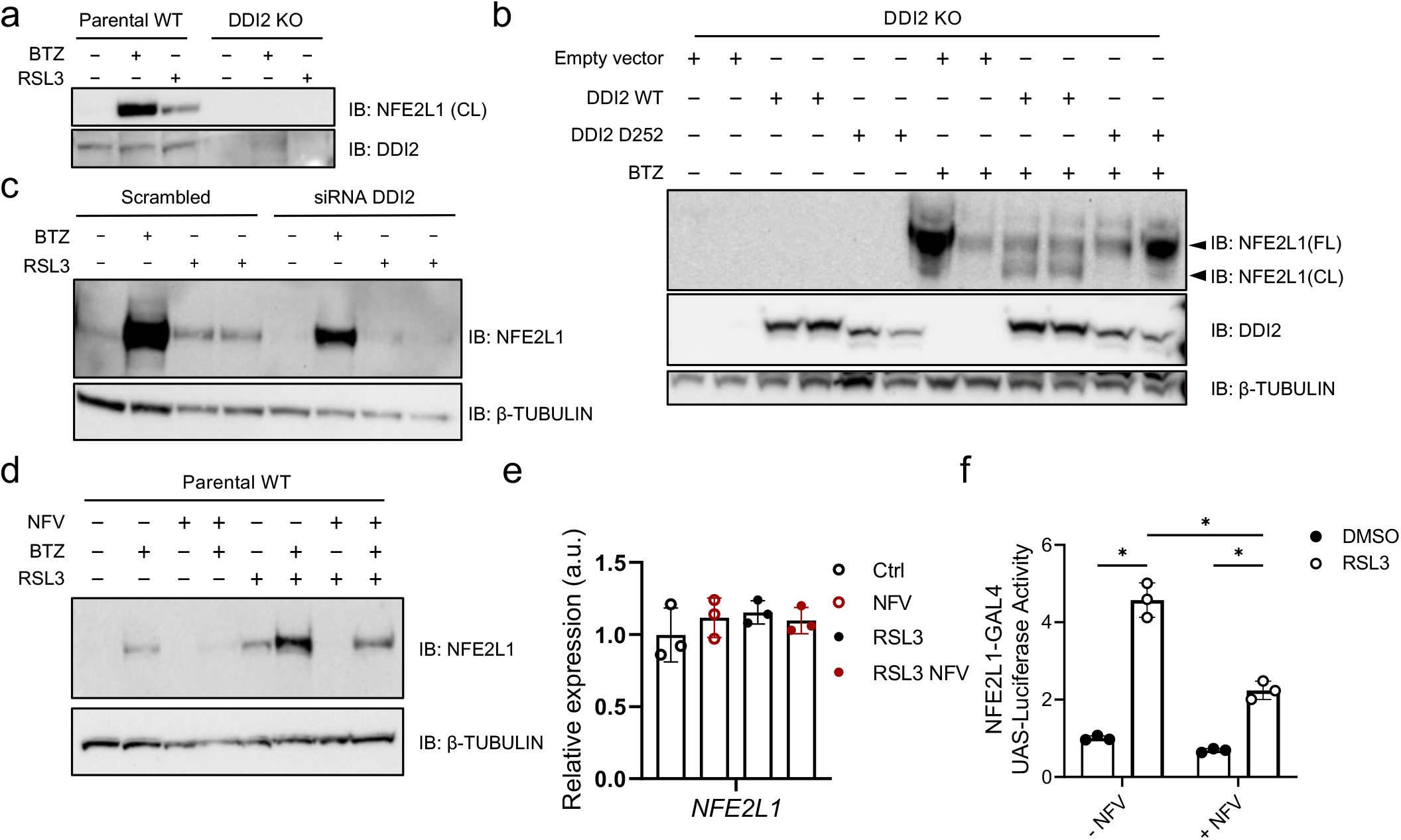
Ddi2 is required for the RSL3-induced proteolytic processing of NFE2L1. (a) Immunoblot of NFE2L1 and DDI2 in EA.hy926 wild-type (Parental WT) and DDI2 knockout (KO) cells treated with 5 μM RSL3 and 100 nM bortezomib (BTZ) for 9 h and 3 h, respectively. (b) Immunoblot of NFE2L1 in DDI2 KO cells transfected with empty vector, wild-type and D252 protease-dead DDI2 carrying plasmids followed by 3 h treatment with 100 nM BTZ (FL: full-length form ca. 120 kDa; CL: cleaved ca. 95 kDa). (c) Immunoblot of NFE2L1 and DDI2 in WT EA.hy926 cells treated with DDI2 siRNA and 3 h of 100 nM BTZ. (d) Immunoblot of NFE2L1 in WT EA.hy926 cells after treatment with 50 μM of protease DDI2 inhibitor nelfinavir (NFV) for 4 h, 5 μM RSL3 for 9 h, and 100 nM BTZ for 3 h. (e) Relative gene expression of NFE2L1 in cells treated with 10 μM of NFV and 5 μM RSL3 for 9 h. f) NFE2L1 luciferase nuclear translocation reporter assay in HEK293a cells after 24 h treatment with 5 μM of NFV and 20 h with 5 μM RSL3.

Silencing DDI2 expression by transfection of siRNA also showed lower levels of the cleaved form of NFE2L1 in cells treated with BTZ and RSL3 (Fig. 2c). Next, we used the established chemical inhibitor of DDI2 nelfinavir (NFV), which is clinically used for the treatment of patients with HIV infections (24). Both BTZ and RSL3-induced activation of NFE2L1 was lower in cells treated with NFV compared to control treated cells (Fig. 2d). This was independent of *NFE2L1* gene expression, which was unchanged (Fig. 2e). We used the chemical inhibition strategy in a nuclear translocation reporter assay, which is a surrogate for NFE2L1 transcriptional activity (20). NFV treatment led to lower reporter activity when cells were stimulated with RSL3, indicating that the nuclear translocation of NFE2L1 during ferroptosis is enabled by DDI2 (Fig. 2e). In summary, these experiments show that DDI2 facilitates the RSL3-induced proteolytic activation and nuclear translocation of NFE2L1.

### DDI2 is required for the UPS bounce back response during ferroptosis

To analyze the impact of the DDI2-NFE2L1 switch for maintaining proper proteostasis during ferroptosis, we analyzed ubiquitylation levels of DDI2 KO compared to WT cells. Immunoblot of BTZ showed an accumulation of ubiquitylated proteins of high molecular weights in DDI2 KO cells (Fig. 3a). Interestingly, this effect on ubiquitin levels was also observed when the cells were treated with RSL3, highlighting the necessity of DDI2 for maintaining proper proteostasis during ferroptosis (Fig. 3a). In-gel measurement of proteasome activity further showed the importance of DDI2 in regulating baseline proteasome activity, as cells treated with NFV showed both lower 30S activity and expression of proteasome subunits in the native PAGE (Fig. 3b). To check if proteasome activity in the context of ferroptosis was also affected by inhibiting DDI2 we used fluorometric peptides and measured the proteasome activity, which revealed that trypsin-like subunit activity of the proteasome was lower upon loss of DDI2 (Fig. 3c). This shows that DDI2 is required for sustaining proteasomal activity during ferroptosis. While GO analysis indicated that proteasome-mediated ubiquitin-dependent protein catabolic process is among the most impacted pathways (Fig. 3d), Volcano plot visualization and principal component analysis of the ubiquitome data further proved higher ubiquitylation status in DDI2 KO cells compared to parental WT cells (Fig. 3e). In summary, the activity of DDI2 is required to activate the NFE2L1-bounce back mechanism of proteasome function in response to ferroptosis initiation to maintain proper proteostasis.

**Fig. 3:**
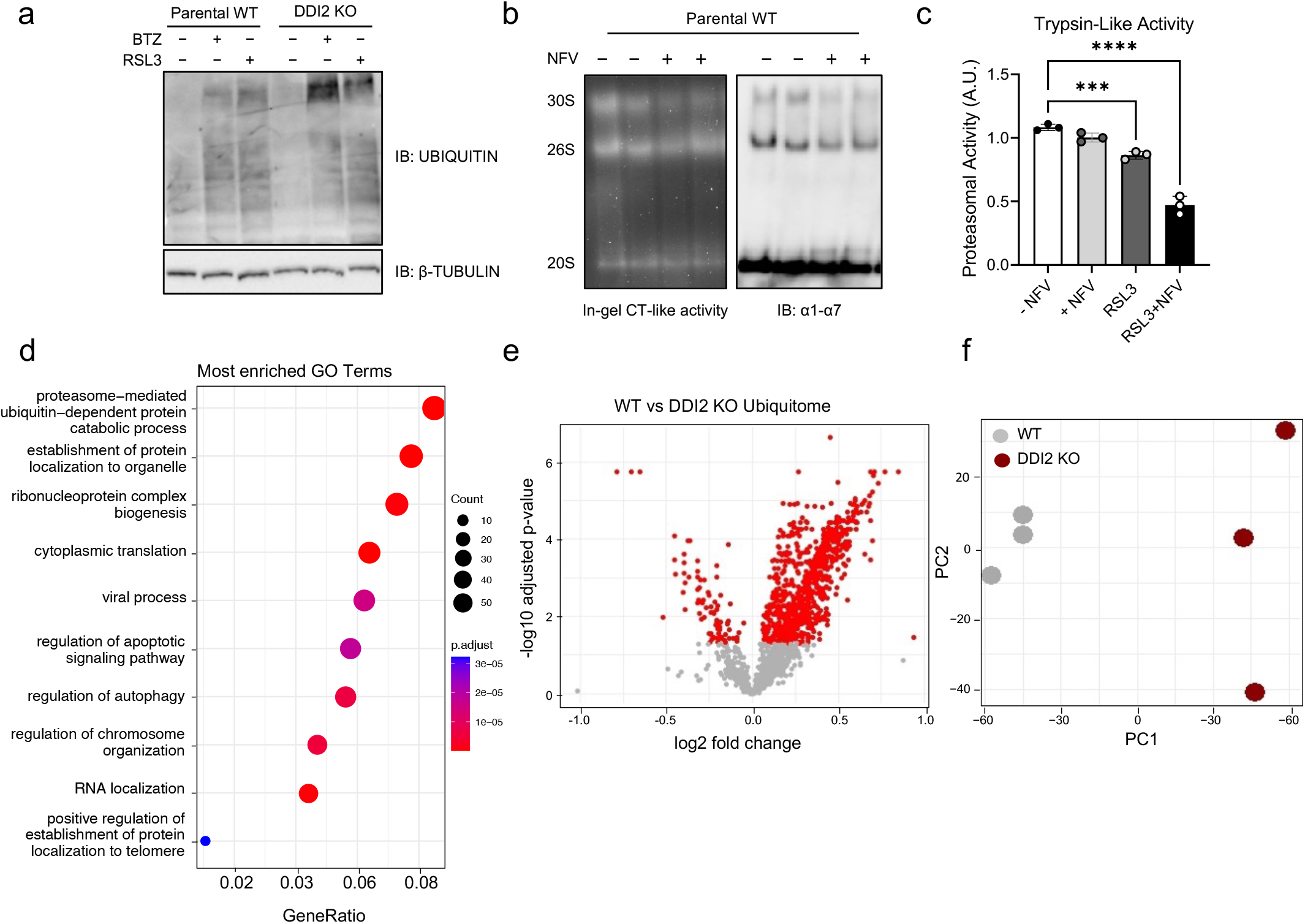
DDI2 is required for the UPS bounce-back response during ferroptosis. (a) Immunoblot of ubiquitin in EA.hy926 cells treated with 5 μM RSL3 and 100 nM bortezomib (BTZ) for 9 h and 3 h, respectively. (b) Native PAGE of EA.hy926 cells treated with 10 μM NFV for 20 h and immunoblot of the α1-7 (20S) subunits. (c) Proteasomal activity of EA.hy926 cells treated with 5 μM NFV for 9 h and 3 h of 5 μM RSL3. (d) Top 10 enriched gene ontology (GO) terms in the ubiquitome of DDI2 (knockout) KO compared to wild-type (WT) parental cells. (e) Volcano Plot of the DDI2 KO cells compared to WT cells *P*_adj_ < 0.05 indicated in red. (n = 3 replicates). (f) Principal component analysis (PCA) of the ubiquitome of EaHy926 WT vs. DDI2 KO cells.

### Proteolytic activation of NFE2L1 by DDI2 protects from ferroptotic cell death

As the main outcome of ferroptosis induction is cell death, we determined the viability of cells upon RSL3 treatment in the presence or absence of the established ferroptosis inhibitor ferrostatin (Fig. 4a). Moreover, we checked if targeting the activity DDI2 with NFV sensitized cells to ferroptosis and potentiated RSL3-induced cell death. NFV treatment increased the cytotoxity of RSL3, which was rescued by ferrostatin (Fig. 4a). We also proved that the appearance of cleaved NFE2L1 protein was dependent on ferroptosis per se as ferrostatin abolished the effect (Fig. 4b). In line with the impact on the NFE2L1-ubiquitin-proteasome system, DDI2 KO cells were more susceptible to ferroptotic death compared to the WT cells (Fig. 4c). We also checked the effect of silencing DDI2 using siRNA on the viability of primary fibroblasts from a patient with Sedaghatian-type Spondylometaphyseal Dysplasia (SSMD), carrying mutant GPX4 compared to healthy control cells. In this distinct genetic ferroptosis cell system, both *NFE2L1* and *DDI2* silencing lowered the viability of cells when ferrostatin was removed. (Fig. 4d). As previously described (13), following the translocation of NFE2L1 to the cytosolic face of ER membrane, NGLY1 removes *N*-linked glycans from NFE2L1 which is a distinct and important posttranslational step for NFE2L1 activation during ferroptosis (Fig. 4d). As others have suggested that deglycosylation of NFE2L1 by NGLY1 takes place prior to cleavage by DDI2 (Fig. 4e), we determined if the NFE2L1-8ND mutant, which contains mutations in the glycosylation sites and is considered constitutively active (13), was processed in the absence of DDI2 under ferroptotic conditions. We transfected WT NFE2L1 and mutant NFE2L1-8ND in DDI2 KO cells. While control cells showed the BTZ- and RSL3-induced cleaved form of NFE2L1, immunoblot of DDI2 KO cells showed no cleaved form of NFE2L1 compared with control cells and this effect was not overcome by using the 8ND mutant form of NFE2L1 (Fig. 4e). Also, to validate if NFE2L1-8ND protected cells from ferroptosis, we performed a cell viability assay, but found no differences between wild-type and mutant forms of NFE2L1 upon RSL3 treatment (Fig. 4f). These experiments highlight the critical role of DDI2 for NFE2L1 activation and subsequent protection from ferroptosis.

**Fig. 4:**
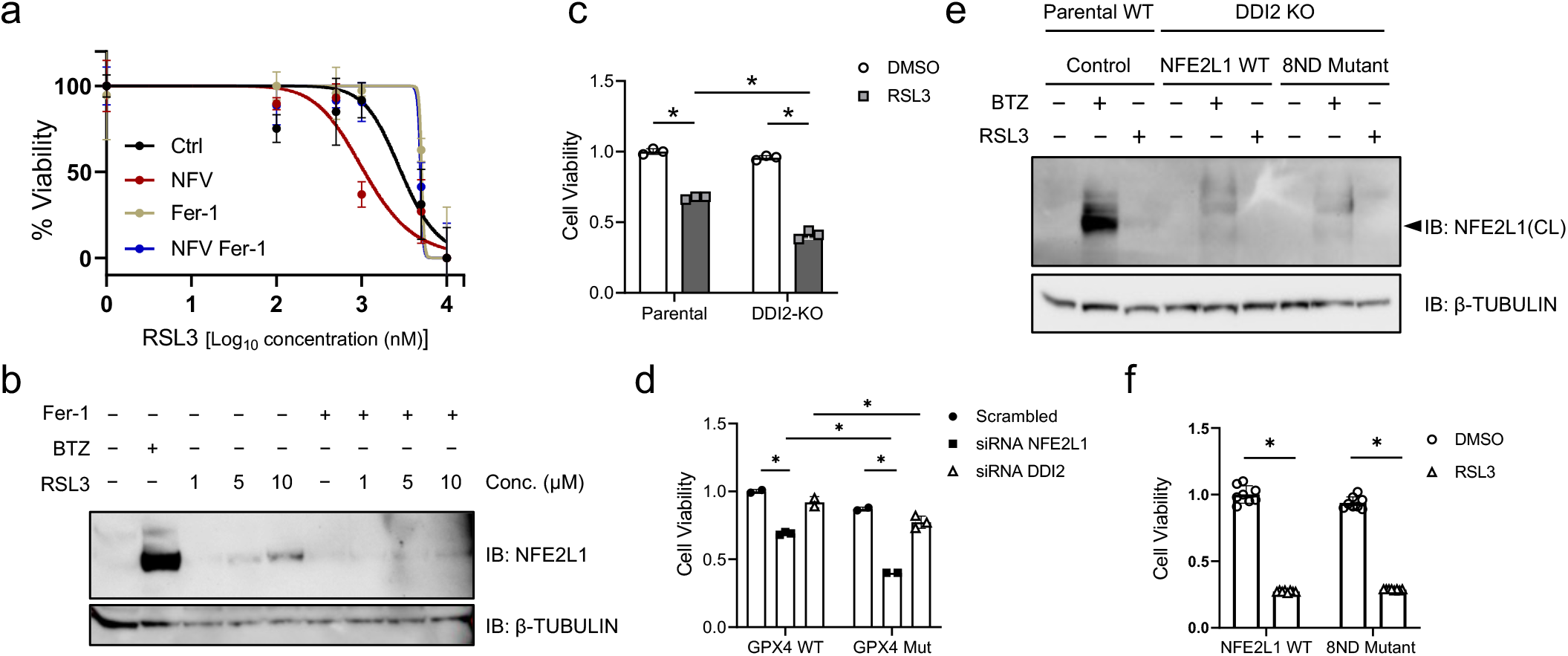
Loss of DDI2 sensitizes cells to chemical and genetic ferroptosis. (a) Cell viability in EA.hy926 cells treated with 5 μM NFV and 10 μM ferrostatin for 20 h. (b) Immunoblot of NFE2L1 in wild-type (Parental WT) and DDI2 knockout (KO) EA.hy926 cells treated 9 h with 10 μM Ferrostatin (Fer-1) and indicated concentrations of RSL3, and 3 h with 100 nM bortezomib (BTZ). (c) Cell viability of DDI2 KO versus WT EA.hy926 cells after 20 h of treatment with 5 μM RSL3. (d) Cell viability of SSMD cells (GPX4 mut) and healthy controls (GPX4 WT) after siRNA-mediated silencing of *NFE2L1* and *DDI2*. (e) Immunoblot of 95 kD cleaved form of NFE2L1 in WT and DDI2 KO EA.hy926 cells transfected with WT NFE2L1 and NFE2L1-8ND mutant plasmids followed by 9 h treatment with 5 μM RSL3 and 100 nM BTZ. (f) Cell viability of DDI2 KO cells transfected with WT NFE2L1 and NFE2L1-8ND mutant plasmids followed by 9 h treatment with 5 μM RSL3.

## Discussion

The execution of ferroptosis and molecular approaches to manipulate its outcomes is an emerging field of research with important implications for metabolism, cancer, and neurodegenerative diseases. While the UPS is clearly involved in ferroptosis (9), little was known about the nature and relevance of this observation. Here, we unequivocally demonstrate the RSL3-induced recalibration of UPS and activation of the proteasome bounce-back mechanism of NFE2L1 by DDI2. In the absence of DDI2, cells accumulate hyperubiquitylated proteins, reflecting a maladaptive regulation of the UPS. Consequently, DDI2 is required to proteolytically activate NFE2L1 and protect from RSL3-induced and genetically driven ferroptotic cell death.

Upon induction of ferroptosis, cells are required to increase proteostatic defense mechanisms, possibly caused by lipid ROS-induced perturbations before the eventual death of the cell. Usually, the proteasome defuses and recycles damaged or obsolete proteins by ubiquitin-dependent and independent degradation. Perhaps, one key finding of our study is that RSL3-induced ferroptosis compromises proteasome activity. However, the nature of this insult was distinct from chemical proteasome inhibition by itself, arguing for more than impact on the ubiquitome. In other words, the recalibration of UPS and ubiquitome during ferroptosis is not only caused by proteasome inhibition. Interestingly, among the ubiquitylated proteins increased upon RSL3 treatment we found many UPS components, including those constituting proteasomes itself. Future work will have to investigate the relevance of these modifications and delineate the posttranslational regulation of proteasome activity during ferroptosis.

In our global ubiquitome analysis, we found several known players that either suppress or promote ferroptosis. Previous studies investigating deubiquitylating enzymes have found a role for ubiquitylation and degradation of GPX4, Nedd4 or VDAC2/3 (25, 26) but the relevance for these mechanisms is still unclear. We also find GPX4 and enzymes of glutathione production to be hyperubiquitylated upon RSL3 treatment and in the absence of DDI2, suggesting that there is a specific impact on reduced proteasome function on key pathways of ferroptosis. On a more general note, our results show lack or dysfunction of DDI2 lead to a global increase in ubiquitylation and reduced survival of the cells when ferroptosis is induced. While hyperubiquitylation increases proteotoxic stress levels, it is possible that the turnover of certain key regulators is disturbed and contributes to the observed ferroptosis sensitivity of DDI2 KO cells. However, as our means of inducing ferroptosis are largely GPX4-dependent, GPX4 is likely not required for the execution of this response. We have previously shown that also other distinct means of inducing ferroptosis such as erastin or FIN56 exhibit changes in proteasomal activity (9). Thus, reduced proteasome function is a hallmark of ferroptosis, and there is overlap between BTZ and RSL3-induced hyperubiquitylation.

The role of DDI2 in the complex series of events leading to NFE2L1 activation should be viewed as required and permissive for the bounce-back response. While reconstituting DDI2 KO cells with WT DDI2 rescued RSL3-induced NFE2L1 activation, a protease-deficient mutant did not. However, proteolytic processing is not the only posttranslational modification required for the activation of NFE2L1, as NGLY1 plays a critical role deglycosylation of NFE2L1 and modulates susceptibility of cells to ferroptosis (13). In our hands, overexpression of deglycosylated NFE2L1 did not rescue the sensitivity of DDI2 KO cells, which underlines the importance of the DDI2-mediated proteolytic cleavage step for the activation of NFE2L1 during ferroptosis. Our data using NFV for inhibiting DDI2 indicate that this could be a potential add-on treatment for targeting ferroptosis in cancer therapy, similar to what has been rationalized for proteasome inhibitors. Interfering with proteasome function and preventing NFE2L1 activation at the same time might further sensitize cancer cells to ferroptosis. Finally, NFE2L1 is also ubiquitylated by the E3 ubiquitin ligase Hrd1 at the ER for ER-associated protein degradation (14), by β-TrCP in the nucleus for removal of NFE2L1 (27), or by UBE4A for cleavage by DDI2 (28), but the details of these regulatory steps for ferroptosis remain unexplored. It will be important to further dissect the posttranslational regulation of NFE2L1 during ferroptosis, as this might reveal more angles to therapeutically exploit the calibration of the UPS for manipulating ferroptosis outcomes.

## Acknowledgement

We thank Silvia Weidner and Thomas Pitsch for excellent technical assistance. We are grateful to Joel Guerra and Leonardo Matta for help with the experiments. We thank all lab members for discussions and the enjoyable atmosphere. We thank Scott Dixon for providing us with the NFE2L1-8ND construct. A.O. was supported by the Integrated Research Training Group of the Deutsche Forschungsgemeinschaft (DFG) CRC1123. S.K. was supported by the Ludwig-Maximilians-University Medical Faculty program FöFoLe for MD students. S.M. and E.K. was supported by DFG RTG2719 PRO. D.T.H. and N.K. were supported by DFG Emmy*-*Noether Program (KR5166/1-1). A.B. was supported by DFG (SFB1123-B10 & SPP2306 BA4925/2-1), the Deutsches Zentrum für Herz-Kreislauf-Forschung (DZHK), and the European Research Training Group (ERC) Starting Grant PROTEOFIT. We acknowledge BioRender.com for help with the figures. We apologize to colleagues whose work we could not cite due to space limitations.

## Author contributions

A.O. designed and performed experiments, analyzed data, and wrote the manuscript. S.K., I.L.L., D.H., and N.K. performed and analyzed proteomics. N.W. and B.B. helped with cell analysis. S.M., S.H.-M., and E.K. generated cell models. A.B. designed experiments, analyzed data, and wrote the manuscript. All authors read and commented on the manuscript.

## Supplementary figures & tables

**Supplementary Fig. 1:**
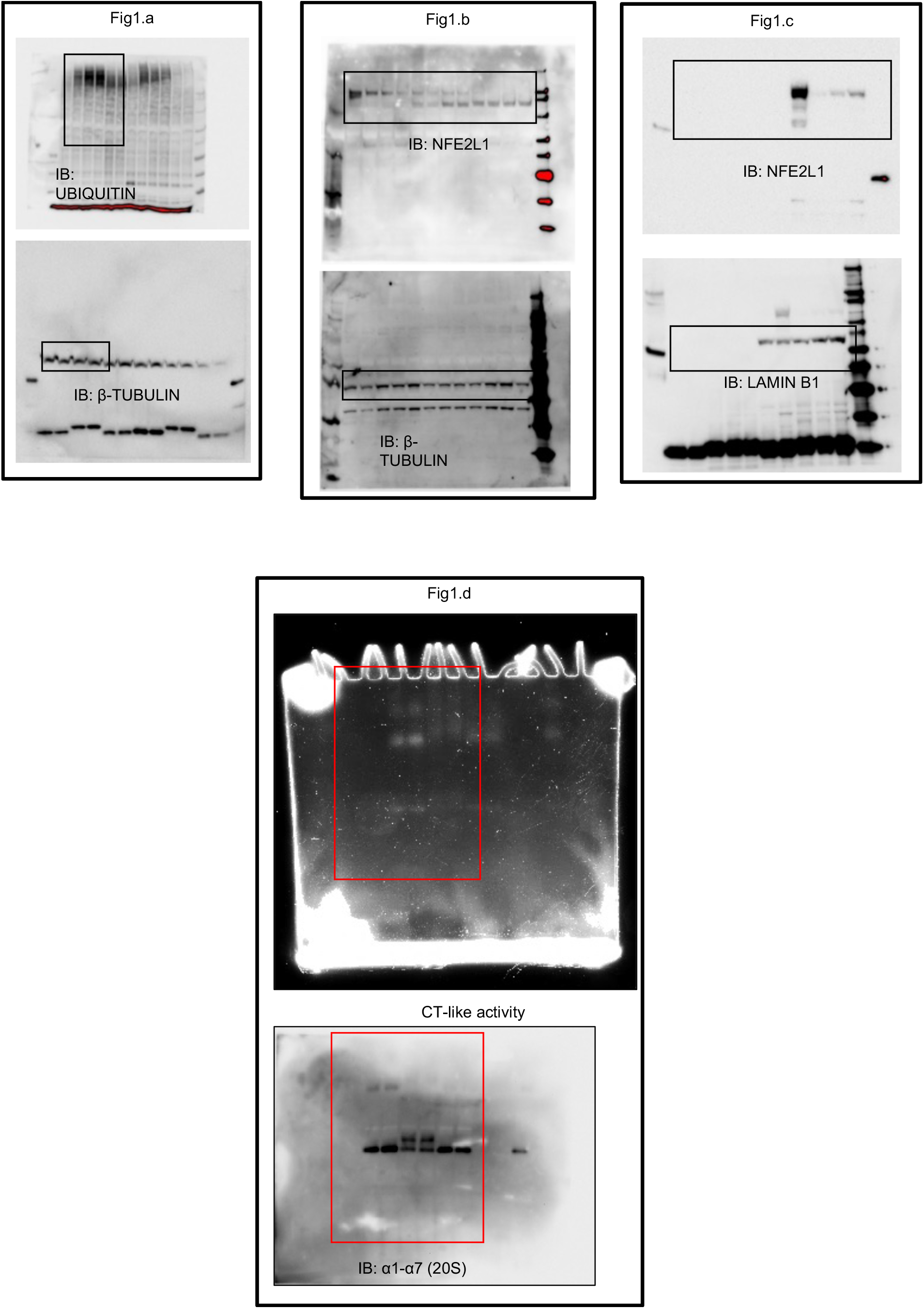
Uncropped immunoblots and gel pictures.

**Supplementary Fig. 2:**
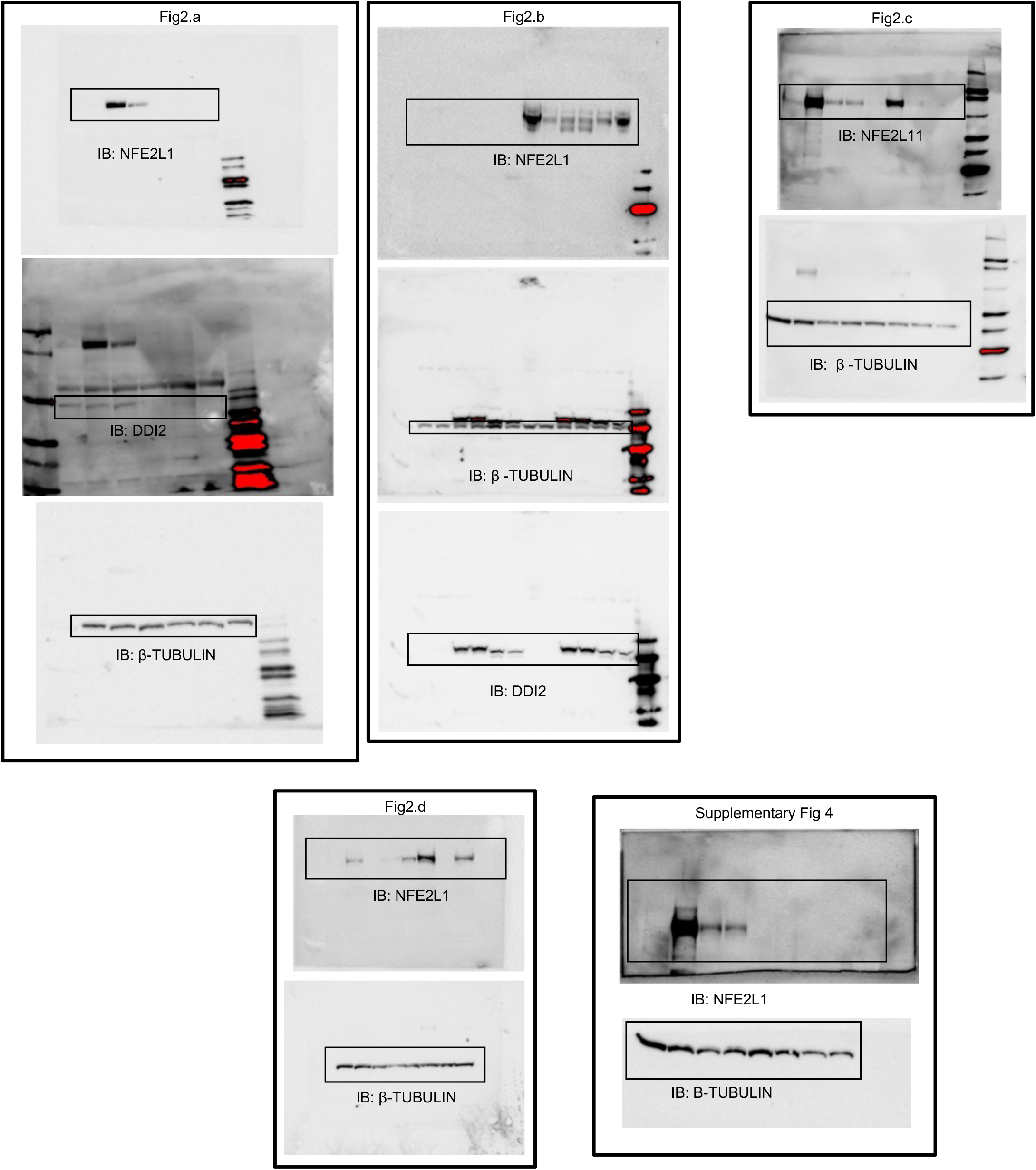
Uncropped immunoblots.

**Supplementary Fig. 3:**
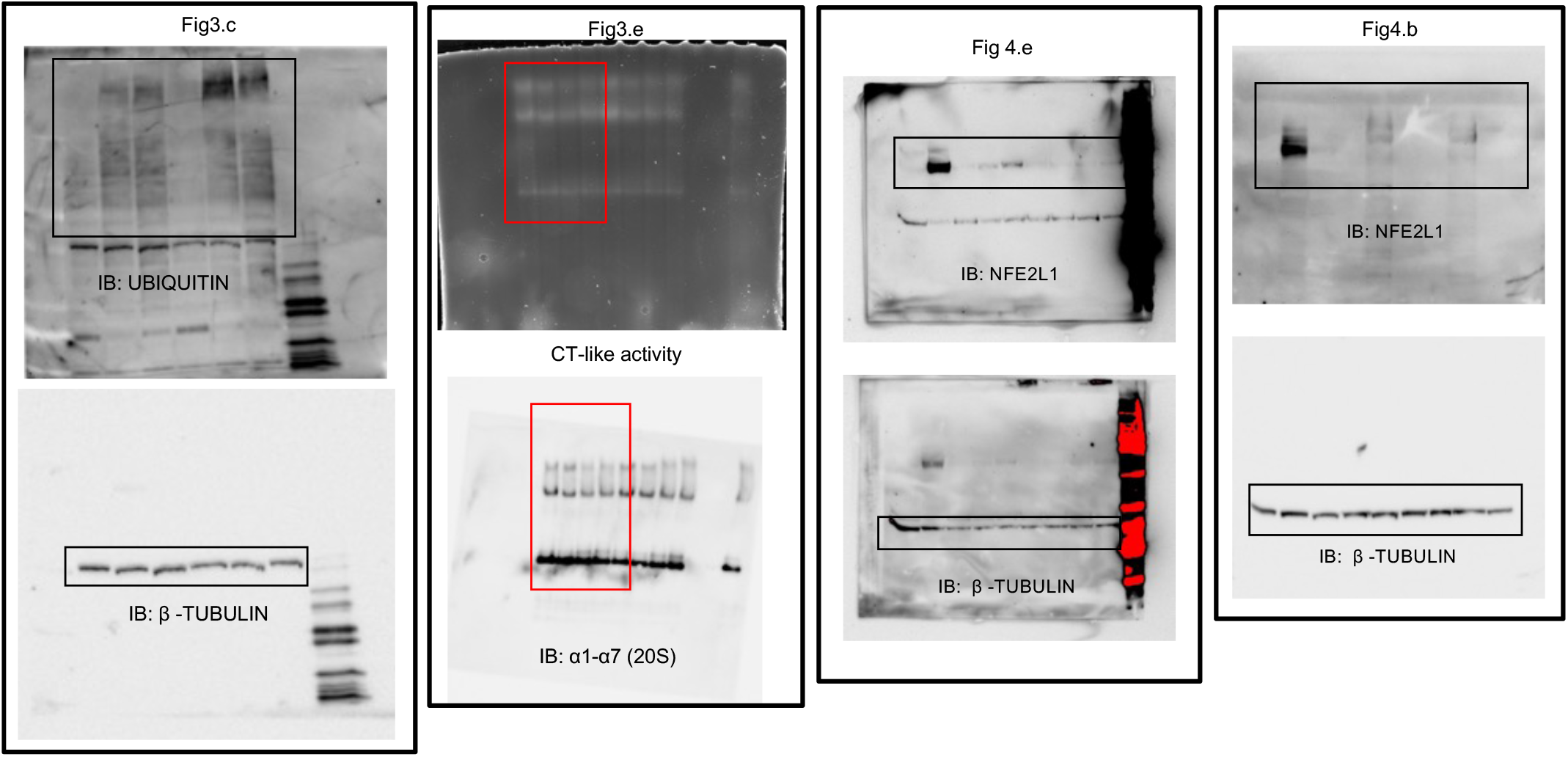
Uncropped immunoblots and gel pictures.

**Supplementary Fig. 4.**
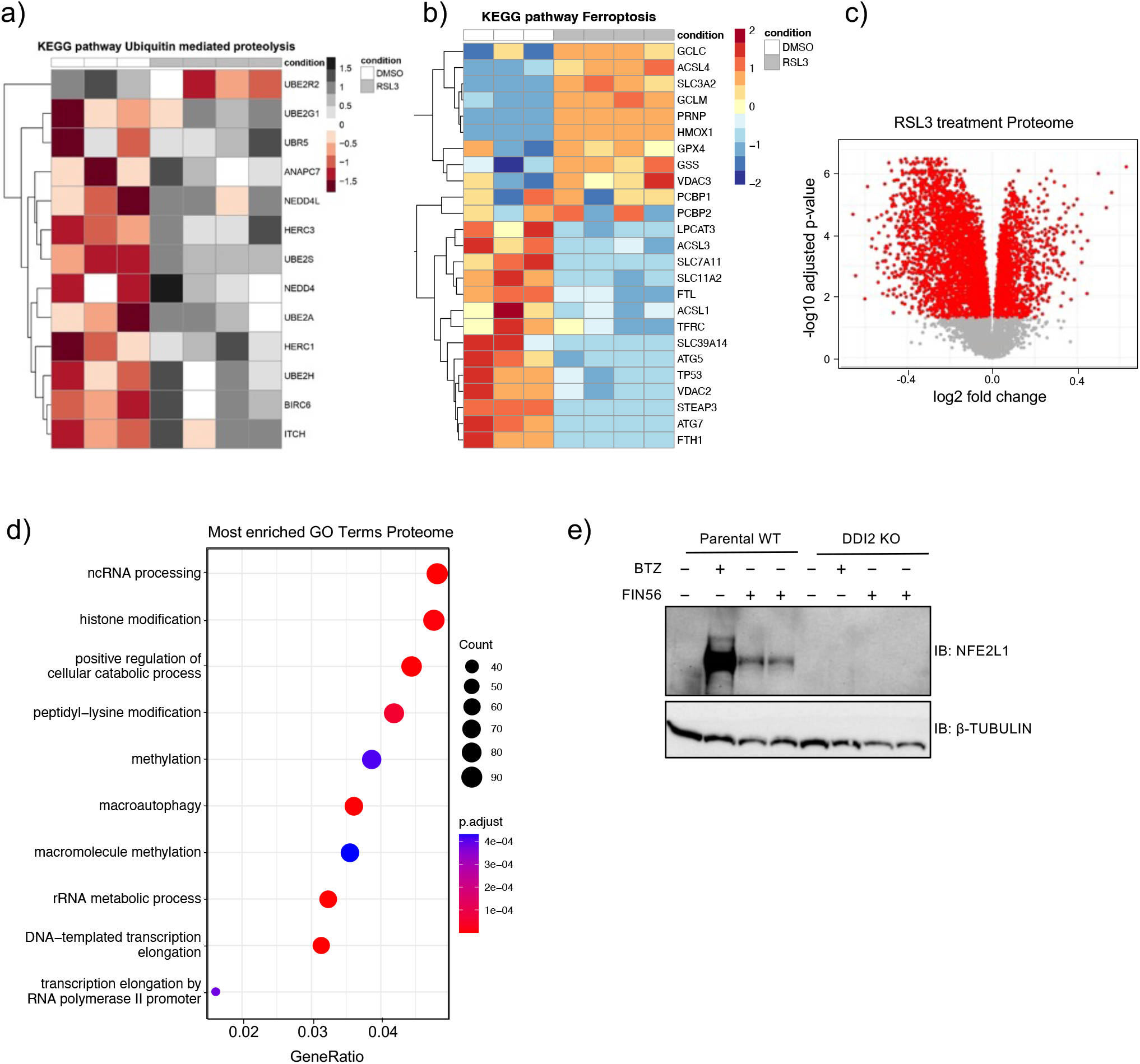
(a) KEGG pathway analysis of ubiquitin mediated proteolysis from ubiquitome of EaHy926 cells treated with 5 μM of RSL3 in the total proteome. (b) KEGG pathway analysis of ferroptosis from total proteome of EaHy926 cells treated with 5 μM of RSL3. (c) Volcano Plot of the proteome of EaHy926 cells treated for 9 h with 5 μM RSL3 *P*_adj_ < 0.05 indicated by red color. (n=4 replicates) (d) Top 10 enriched gene ontology (GO) terms in the proteome of EaHy926 cells treated for 9 h with 5 μM RSL3. (e) Immunoblot of NFE2L1 in wild-type EA.hy926 and DDI2 KO cells treated with 10 μM FIN56 for 9 h and 100 nM BTZ for 3 h.

**Supplementary Table 1:**

| Antibody | Company (Cat. No.) | Dilution |
| --- | --- | --- |
| DDI2 | Abcam (ab197081) | 1:1000 |
| NFE2L1 | Cell signaling technologies (D9K6P) | 1:500 |
| β-TUBULIN | Cell signaling technologies (2146) | 1:1000 |
| UBIQUITIN (P4D1) | Cell signaling technologies (3936) | 1:1000 |
| 20S Proteasome $\alpha$ subunit | Abcam (ab22674) | 1:1000 |
| Anti-Lamin B1 | Abcam (ab229025) | 1:1000 |
| Anti-rabbit IgG, HRP-linked | Cell signaling technologies (7074) | 1:10000 |
| Anti-mouse IgG, HRP-linked | Cell signaling technologies (7076) | 1:10000 |

**Supplementary Table 2:**
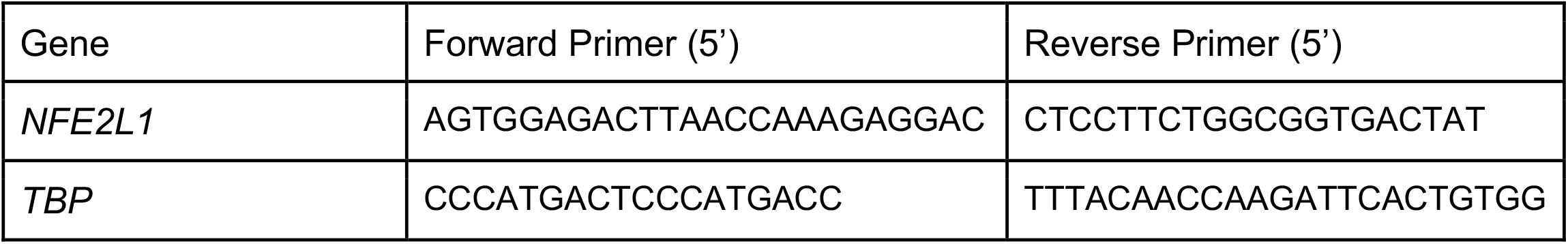

